# Spatially uniform establishment of chromatin accessibility in the early *Drosophila* embryo

**DOI:** 10.1101/195073

**Authors:** Jenna E. Haines, Michael B. Eisen

**Affiliations:** Department of Molecular and Cell Biology, University of California, Berkeley, CA; Department of Integrative Biology, University of California, Berkeley, CA; Howard Hughes Medical Institute, University of California, Berkeley, CA

## Abstract

As the *Drosophila* embryo transitions from the use of maternal RNAs to zygotic transcription, domains of open chromatin, with relatively low nucleosome density and specific histone marks, are established at promoters and enhancers involved in patterned embryonic transcription. However, it remains unclear whether open chromatin is a product of activity - transcription at promoters and patterning transcription factor binding at enhancers - or whether it is established by independent mechanisms. Recent work has implicated the ubiquitously expressed, maternal factor Zelda in this process. To assess the relative contribution of activity in the establishment of chromatin accessibility, we have probed chromatin accessibility across the anterior-posterior axis of early *Drosophila melanogaster* embryos by applying a transposon based assay for chromatin accessibility (ATAC-seq) to anterior and posterior halves of hand-dissected, cellular blastoderm embryos. We find that genome-wide chromatin accessibility is remarkably similar between the two halves. Promoters and enhancers that are active in exclusively one half of the embryo have open chromatin in the other half, demonstrating that chromatin accessibility is not a direct result of activity. However, there is a small skew at enhancers that drive transcription exclusively in either the anterior or posterior half of the embryo, with greater accessibility in the region of activity. Taken together these data support a model in which regions of chromatin accessibility are defined and established by ubiquitous factors, and fine-tuned subsequently by activity.

## Introduction

During early embryogenesis all animal genomes undergo a transition from a largely quiescent to a highly active state with widespread zygotic transcription [1]. This process, known as the maternal-to-zygotic transition (MZT), involves a major reorganization of chromatin during which active and inactive regions, distinguished by nucleosome composition, density and posttranslational modifications, are established [2–6]. It is generally thought that active - or “open” - chromatin facilitates the binding of polymerases, transcription factors, and other proteins to target sequences, while inactive - or “closed” - chromatin limits the scope of their activity, although the degree to which chromatin state is instructive remains controversial [7,8]. Two important open questions are how genomic locations of active and inactive chromatin become encoded in the genome and how their activity state is established, especially during early embryogenesis when no pre-formed patterns exist.

In *Drosophila melanogaster,* zygotic transcription largely begins at the seventh syncytial mitotic cycle (although there is evidence for low levels of transcription from the beginning of embryogenesis [9]) which gradually increases until the end of mitotic cycle 13, when the embryo has several thousand nuclei and widespread zygotic transcription is observed [10,11]. Many of the genes activated during the MZT produce mRNAs that have spatially restricted distributions. These patterns are established through the activity of transcriptional enhancers, *cis*-regulatory sequences that integrate activating and repressive inputs from well-characterized, patterning transcription factors to produce novel, increasingly precise transcriptional outputs [12–15].

It is widely assumed that the interactions among patterning factors and the DNA to which they bind determines which sequences will function as enhancers and that their competition with nucleosomes and recruitment of chromatin remodeling factors establish chromatin accessibility at selected sites [16–19]. However, we and others have shown that a parallel system involving the maternally-deposited transcription factor Zelda (ZLD) plays an important role in this process. Zelda binds prior to the MZT to a large fraction of the enhancers and promoters that become active once widespread zygotic transcription begins [20,21]. Most MZT enhancers and promoters contain conserved Zelda binding sites that are highly predictive of both transcription factor activity and chromatin accessibility [21]. Furthermore, Zelda binding is associated with changes to chromatin, including nucleosome depletion and specific post-translational modifications of histones [2,21–24].

Based on these and other observations, we have proposed that the activation of enhancers and promoters in the early embryo is a two-step process, with Zelda determining which regions of the genome will have accessible chromatin, while the binding of patterning transcription factors to these open regions determines their transcriptional output. An important prediction of this model is that, because Zelda is ubiquitously expressed, enhancers and promoters should be accessible across the entire embryo. If, however, patterning factors play a dominant role in establishing accessibility, we expect genomic regions to be in an accessible state only where the appropriate patterning factors are active.

To explore these alternative hypotheses, here we compare chromatin accessibility in anterior and posterior regions of the *D. melanogaster* embryo. We find that genome-wide chromatin accessibility is remarkably similar between the two halves, with promoters and enhancers active exclusively in half of the embryo being equally accessible in both halves. However, we also found that at enhancers that drive expression exclusively in the anterior or posterior half of the embryo, accessibility is distinctly but modestly skewed in the direction of activity, with those with the strongest skew having an anterior bias. These data suggest that chromatin accessibility in the early embryo is generally open at regulatory regions and is only modestly patterned based on activity.

## Results

### Spatially resolved ATAC-seq is robust and consistent with whole embryo measurements of chromatin accessibility

To determine the extent to which chromatin accessibility is spatially patterned along the anteroposterior axis in the early embryo, we manually separated anterior and posterior embryo halves and performed a modified ATAC-seq [25] protocol on each half separately. Briefly, we collected cellular blastoderm embryos (mitotic cycle 14, embryonic stage 5), flash froze them in liquid nitrogen, and then sliced each embryo with a chilled scalpel at the anteroposterior midline, separating anterior and posterior halves into separate pools (Fig. 1a). We isolated nuclei from 20 anterior halves (in duplicate), 20 posterior halves (in duplicate), 10 frozen unsliced embryos, and a mixed sample containing a subset of nuclei from anterior and posterior samples and applied the ATAC-seq “tagmentation” process to each sample. We sequenced the resulting libraries, mapped reads to the *D. melanogaster* genome and normalized the data using standard methods (Fig. S1).

**Figure 1.**
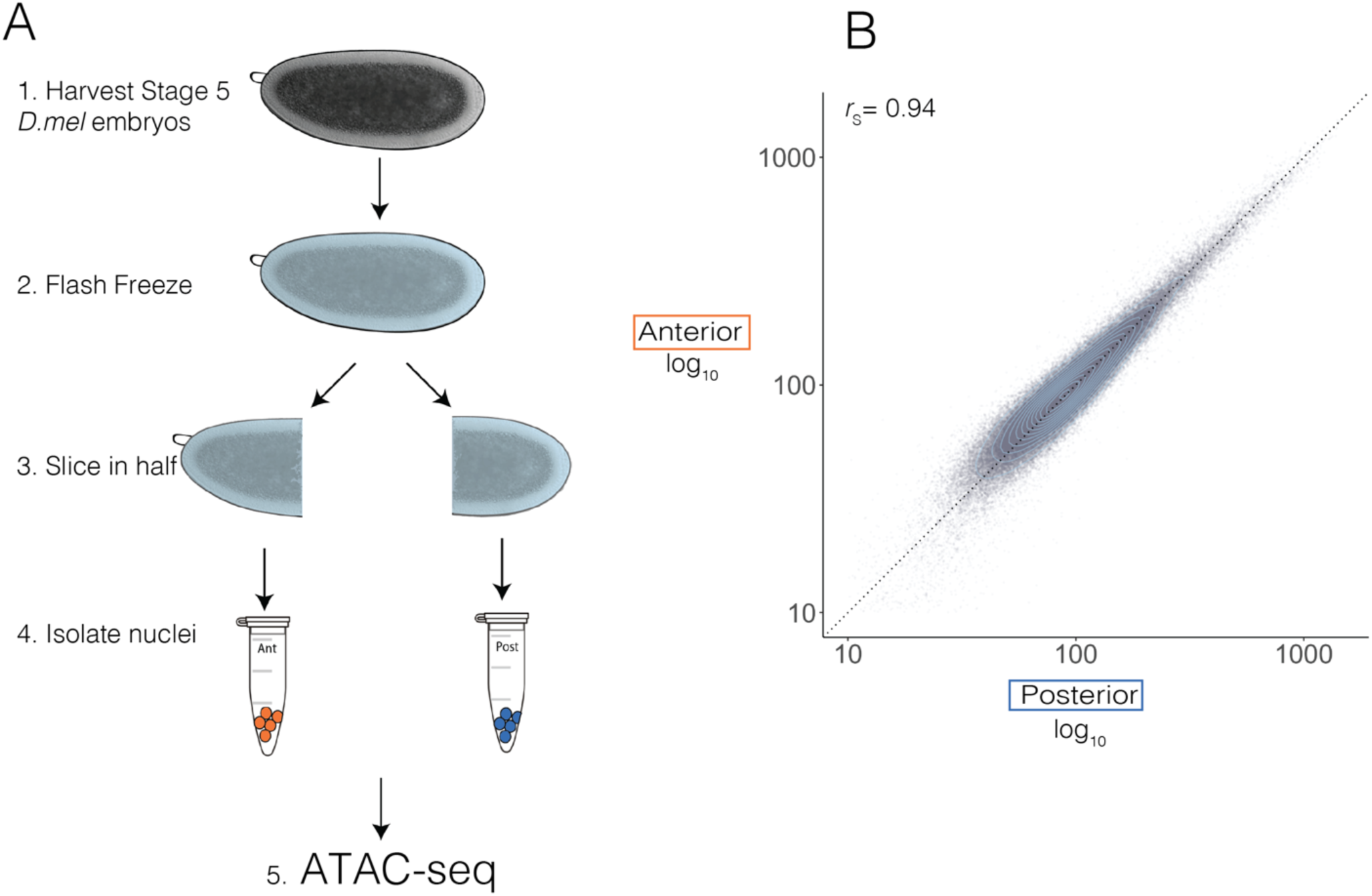
ATAC-seq on dissected, frozen, embryo halves. (A) Stage 5, hand sorted *Drosophila* embryos were flash frozen over dry ice in a buffer containing 5% glycerol and manually sliced in half with a scalpel. Twenty anterior and posterior halves were collected, homogenized, and the nuclei were isolated. ATAC-seq was then performed as described in (Buenrostro et al. 2013) with three times Tn5 transposase. (B) Scatterplot of normalized ATAC- seq signal over 1kb adjacent windows that tile the *Drosophila* genome in posterior (x) and anterior (y) samples shows high degree of correlation between the anterior and posterior halves. The Spearman correlation coefficient (denoted by *r*S) is 0.94. X and Y are log transformed. Light blue circles denote point density.

Data from sliced and unsliced whole embryo samples were nearly identical, demonstrating that the slicing process does not introduce any biases (*r*_S_ > 0.90, Fig. S2a-c). Both halves were highly similar to published DNAseI hypersensitivity data from similar embryo stages, demonstrating that our embryo preparation protocol coupled with ATAC-seq can accurately map accessibility in the equivalent of 10 whole frozen embryos (*r*_S_ > 0.80, Fig. S2a). Biological replicates of anterior and posterior halves that were collected, sliced, and tagmented independently correlate highly with each other (Anterior replicates *r*_S_ = 0.89, Posterior replicates *r*_S_ = 0.82), and were therefore merged for all future analyses (Fig. S2b). We then called peaks on ATAC-seq data from merged anterior and posterior replicates using MACS2 [26].

### Globally similar chromatin accessibility patterns in anterior and posterior embryo halves

Genome-wide, chromatin accessibility in the anterior and posterior halves is remarkably similar (Fig. 1b; *r*_S_ = 0.94 on data binned into 1kb windows), with 73% of all accessibility peaks shared between the two halves of the embryo. The conservation of chromatin accessibility patterns between halves is detailed in Figure 2, which shows the results of our ATAC-seq experiments near loci of three anteroposterior patterned genes (*even-skipped, giant,* and *hunchback*) and one dorsovental patterned gene (*dpp*).

**Figure 2.**
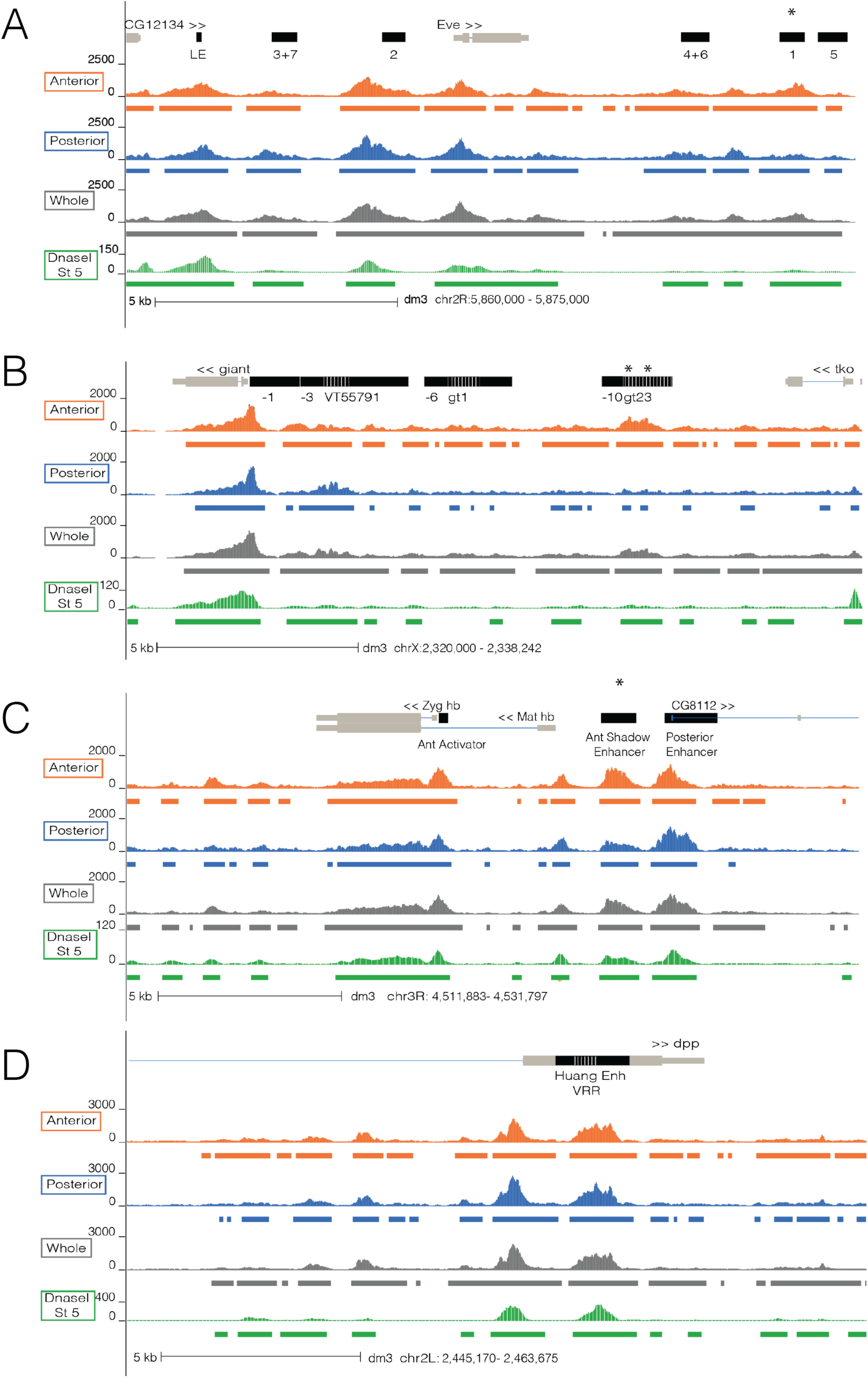
Chromatin accessibility differences and similarities at A-P and D-V patterned loci. Normalized ATAC-seq signal of anterior (orange), posterior (blue), whole embryo (gray) is depicted at the *even-skipped* (A), *giant* (B), and *hunchback* (C) loci which contain A-P patterned enhancers and promoters and at *decapentaplegic* (D), a D-V patterned gene. Chromatin accessibility data derived from DNaseI for stage 5 *Drosophila* embryos is depicted in green (Thomas et al. 2011). Colored bars represent peaks called in anterior (orange), posterior (blue), whole (gray), and in DNaseI data (green). Asterisks denote annotated features that show strong changes in accessibility between the anterior and posterior halves. Light gray brown bars denote the gene annotation while the black bars denote enhancers. Dashed white lines signify two overlapping regions between enhancers.

Each of these anteroposterior loci contains enhancers that are active exclusively in the anterior or posterior half of the embryo (denoted by black bars in Fig. 2). In all cases there are accessibility peaks at these enhancers in both embryo halves. For some, the peaks are of similar heights, such as at *eve* stripes 2 (Anterior ATAC-seq signal / Posterior ATAC-seq Signal - 528/563) and 5 (173/183), *gt* anterior (-1 construct, 388/351), and *gt* posterior (VT55791, 213/246) enhancers. However, there are some examples where accessibility is clearly reduced in the inactive half, such as at *eve* stripe 1 (670/320), the *gt* anterior enhancers 23 (467/189) and - 10 (384/173), and the *hb* anterior “shadow” enhancer (543/298) (Fig. 2, marked by asterisks). Notably, peaks from both halves overlap all A-P patterned enhancers we analyzed with the exception of three anteriorly patterned enhancers (slp1 A, HC_52, and CG9571 fd19B).

### Anteroposterior enhancers are ubiquitously open but accessibility levels track activity

To get a more systematic view of the relationship between transcriptional activity and spatial patterns of chromatin accessibility, we used available genome annotation and functional data to systematically identify A-P and D-V (as a control) patterned enhancers whose transcriptional outputs are restricted to one half of the embryo [27–38] (File S1). We excluded enhancers that did not overlap peaks called in any of the anterior, posterior, or whole samples leaving 98 A-P and D-V patterned enhancers.

Although virtually all of the A-P patterned enhancers we looked at are accessible in both halves, they clearly trend towards greater accessibility in the embryo half where they are active (Fig. 3). Normalized ATAC-seq signal at anterior and posterior enhancers (anterior *r*_S_ = 0.82; posterior *r*_S_ = 0.85) show weaker correlation than all 1kb regions genome-wide (gray; *r*_S_ = 0.94) or D-V patterned enhancers (*r*_S_ = 0.97, Anterior n = 36, orange; Posterior n = 18, blue; Dorsal n = 17, purple; Ventral n = 27, green; Fig. 3a-b).

**Figure 3.**
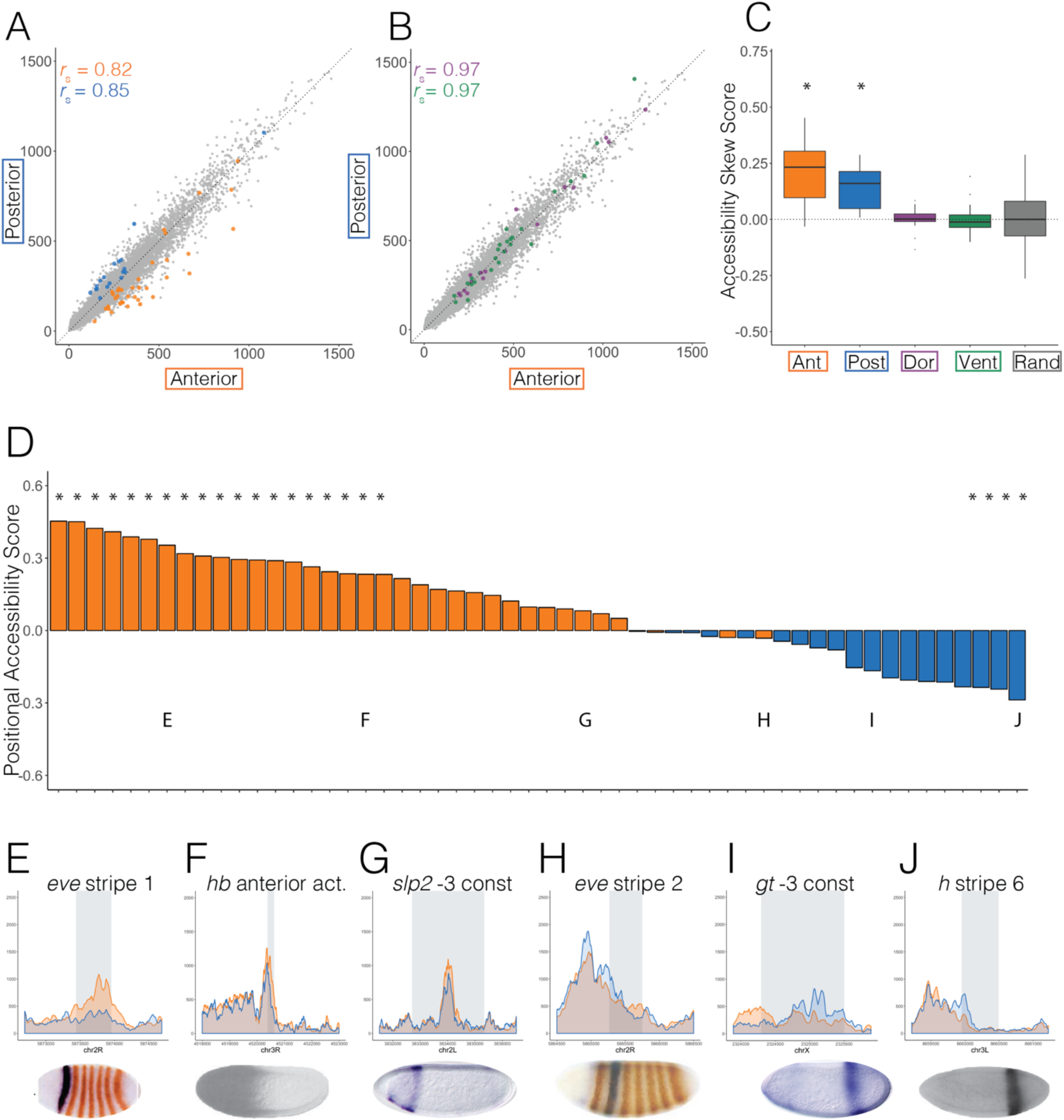
A-P patterned enhancers tend to be more accessible where they are active. Scatterplots showing normalized ATAC-seq signal in anterior (x axis) and posterior (y axis) halves at (A) anterior (orange) and posterior (blue) and (B) dorsal (purple) and ventral (green) patterned enhancers active in Stage 5 embryos and at 1kb adjacent windows tiling the genome (A and B, gray). (C) Boxplots showing the difference in mean and variation between overall accessibility skew scores (methods) for anterior (orange), posterior (blue), dorsal (purple), and ventral (green) enhancers. Accessibility skew scores at random genomic regions (excluding genes and enhancers) with the same number and distribution of total ATAC-seq signal as the patterned enhancer and promoter set depicted here and in Fig. 4 (methods). ANOVA analysis confirms that the means of both anterior and posterior patterned enhancers are significantly different than dorsal and ventral patterned enhancers and selected random regions. (D) Bargraph shows the positional accessibility skew scores (methods) calculated for all anterior (orange) and posterior (blue) patterned enhancers in the dataset (File S1). Asterisks denote enhancers whose accessibility skew scores show statistical significance over random regions (p < 0.05). (E-J) Normalized ATAC-seq signal across 2kb windows centered around eve stripe 1 (E), the *hunchback* anterior activator (F), *slp2* -3 construct (G), *eve* stripe 2 (H), *giant* -3 construct (I), and *hairy* stripe 6 (J) with anterior signal in orange and posterior signal in blue. Gray rectangles denote the location and size of the enhancer. Published *in situ* hybridization images depicting gene expression patterns driven by each enhancer are below each graph (Fujioka et al. 1999; Driever et al. 1989; Schroeder et al. 2004; HÄder et al. 1998).

We next computed a measure of differential accessibility (accessibility skew score) for each enhancer by dividing the difference in accessibility in the active and inactive half by total accessibility, such that positive scores denote loci that are more accessible in the active half, negative scores signify loci that are more accessible in the inactive half, and loci with a score of zero have no difference in accessibility (methods). We found that both anterior and posterior enhancers have a significantly greater mean accessibility skew score than D-V enhancers or random genomic regions with similar accessibility (p_ant_ < 7.8 x 10^-10^ and p_post_ < 0.0016, Fig. 3c and Table S1).

Figure 3d shows positional accessibility skew scores at all A-P patterned enhancers that we analyzed. Strikingly, accessibility at almost all anterior enhancers is skewed towards the anterior while that of posterior enhancers is skewed towards the posterior. This pattern is in contrast to D-V patterned enhancers and promoters (Fig. S3) and A-P patterned promoters (Fig. 4d).

**Figure 4.**
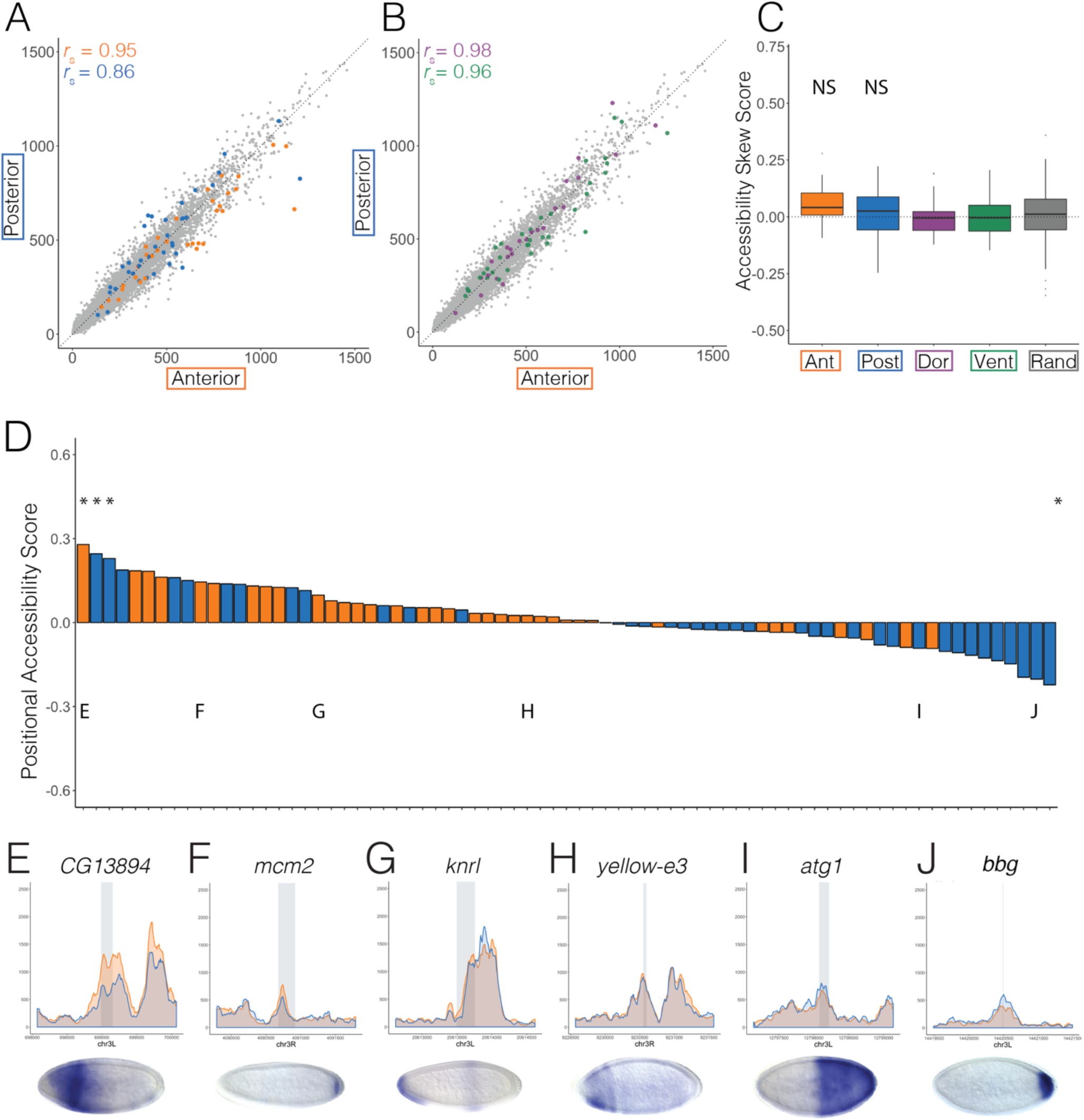
A-P patterned gene promoter accessibility does not correlate with activity. Scatterplots showing normalized ATAC-seq signal in anterior (x axis) and posterior (y axis) halves at (A) anterior (orange) and posterior (blue) and (B) dorsal (purple) and ventral (green) patterned promoters active in Stage 5 embryos and at 1kb adjacent windows tiling the genome (A and B, gray). (C) Boxplots showing the difference in mean and variation between overall accessibility skew scores (methods) for anterior (orange), posterior (blue), dorsal (purple), and ventral (green) promoters. Accessibility skew scores at random genomic regions (excluding genes and enhancers) with the same number and distribution of total ATAC-seq signal as the patterned enhancer and promoter set depicted here and in Fig. 3 (methods). ANOVA analysis confirms that the means of anterior and posterior patterned promoters are not significantly different than dorsal and ventral patterned enhancers and selected random regions. (D) Bargraph shows the positional accessibility skew scores (methods) calculated for all anterior (orange) and posterior (blue) patterned promoters in the dataset (File S1). Asterisks denote enhancers whose accessibility skew scores show statistical significance over random regions (p < 0.05). (E-J) Normalized ATAC-seq signal across 2kb windows centered around *CG13894* (E), *mcm2* (F), *knrl* (G), *yellow-e3* (H), *atg1* (I), and *bbg* (J) with anterior signal in orange and posterior signal in blue. Gray denotes the location and size of the promoter. Published *in situ* hybridization images depicting gene expression patterns driven by each promoter are below each graph (Hammonds et al. 2013; Tomancak et al. 2002; Tomancak et al. 2007).

### Promoters of A-P patterned genes are similarly accessible both when active and inactive

We next examined the promoters of A-P patterned genes using expression data from sections of embryos cryosliced along the A-P axis to curate lists of A-P patterned gene promoters [39]. We only included promoters that overlapped accessibility peaks called on our dataset and have patterned expression confirmed by *in situ* hybridization assays (n= 36 anterior promoters, n = 39 posterior promoters, File S1).

Interestingly, accessibility in the active and inactive halves is much more similar at anterior promoters (*r*_S_ = 0.95) than at anterior enhancers (*r*_S_ = 0.82), and is comparable to D-V enhancers and promoters (Fig. 4 a-b, Dorsal promoters, *r*_S_ = 0.98; Ventral promoters, *r*_S_ = 0.96). On the other hand, posterior promoters and enhancers show similar accessibility correlations between the anterior and posterior halves (*r*_S_ = 0.86), though as a group their mean accessibility skew score is not significantly different than D-V patterned promoters or random regions (Fig. 4c and Table S1). Additionally, there is no distinct skew of accessibility in the direction of activity seen at A-P promoters, in contrast to what we observed for A-P enhancers.

### Anterior accessibility is associated with Bicoid binding while similarly accessible regions are enriched for Zelda binding

We then used published ChIP-seq data of A-P patterning factors from stage 5 *Drosophila* embryos to examine binding patterns at similarly and differentially accessible A-P enhancers [40]. We analyzed Bicoid, Caudal, Knirps, Giant, Hunchback, Kruppel, and Zelda binding data, normalized by the mean signal for each factor. A-P enhancers that are more accessible in the anterior (shades of orange) generally seem to be dominated by Bicoid binding with strikingly little binding from other transcription factors, though there are some exceptions (Fig. 5, Fig. S4). Enhancers more accessible in the posterior (shades of blue) generally have high Caudal, Knirps, Giant, and Kruppel binding with strikingly more diversity in factors bound than at anteriorly accessible enhancers. Interestingly, enhancers with similar accessibility in both halves (shades of white) generally have a high diversity of factors binding - including Zelda (Fig. 5b). These patterns reveal that while transcription factor binding clearly does not completely explain differential chromatin accessibility, there are clear differences in factor composition and density between differentially and similarly accessible enhancers.

**Figure 5.**
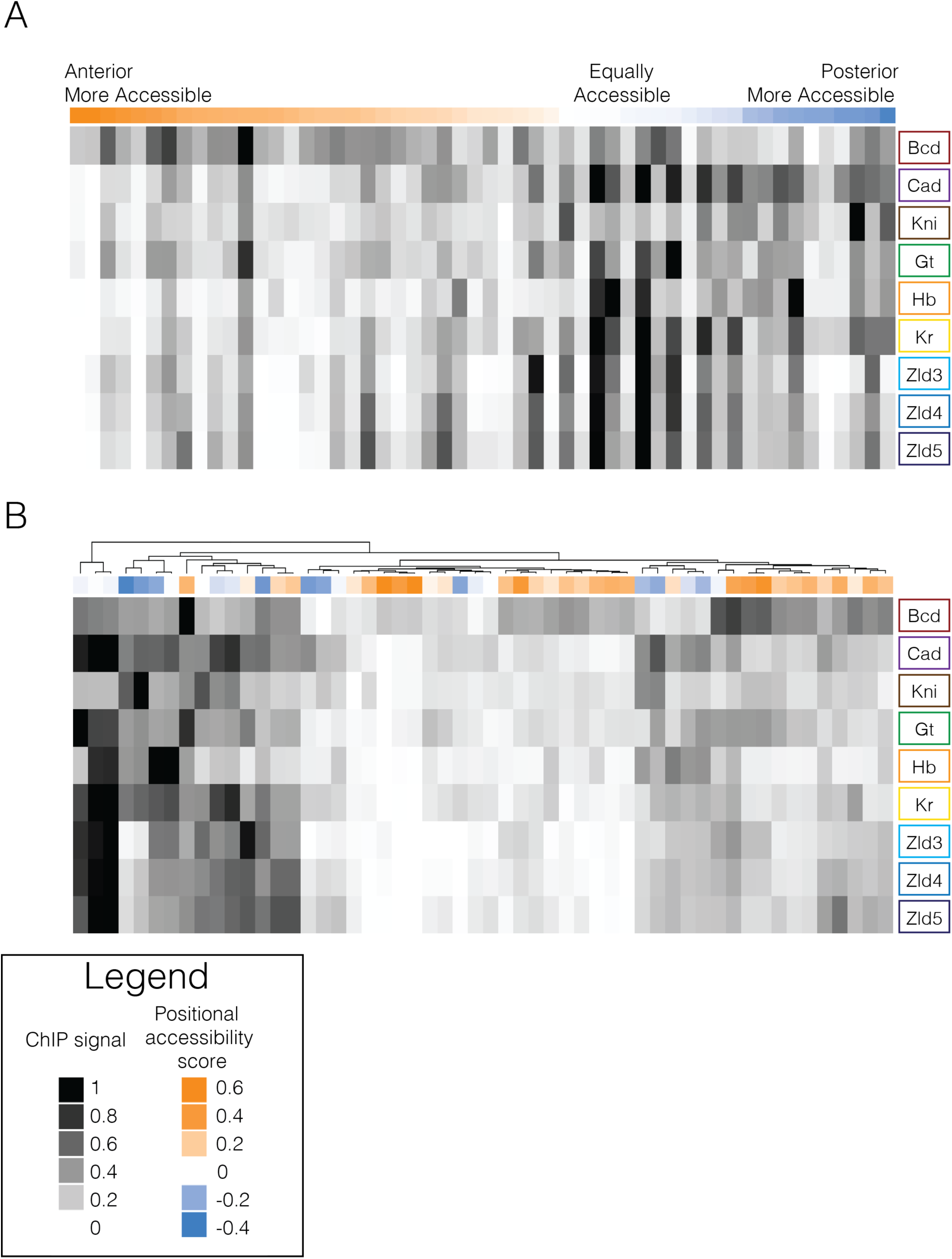
Transcription factor binding and differential accessibility at A-P patterned enhancers. ChIP-seq data for Bicoid, Caudal, Knirps, Giant, Hunchback, Kruppel, and Zelda from three stages (stage 3,4, and 5) from (Bradley et al. 2010; Harrison et al. 2011) normalized to the mean of each factor and scaled between 0 and 1 summed over a 5kb window around each A-P enhancer. White represents the minimum signal and black represents the maximum ChIP signal for that transcription factor. Above each heatmap is a colored bar that represents the positional accessibility skew score for each A-P enhancer with orange representing enhancers that are more accessible in the anterior, blue representing those that are more accessible in the posterior, and white representing those that are not differentially accessible between the two halves. (A) Enhancers are ordered by positional accessibility score. (B) Enhancers are hierarchically clustered using complex heatmap package in R (Gu et al. 2016).

## Discussion

In both mammalian and fly differentiated tissues there is a clear relationship between the activity of a DNA element and chromatin accessibility [17,41–44]. Our data demonstrate that the early *Drosophila* embryo is different, in that regions open in one half of the embryo are almost always open in the other, even when looking at promoters and enhancers of genes that are active exclusively in one half of the embryo.

This finding adds to a growing body of evidence that the chromatin landscape of the early embryo is distinct from that of differentiated tissues found in later stage embryos. A recent study assaying chromatin accessibility in single nuclei of developing *Drosophila melanogaster* embryos found that 2 to 4 hour old embryos cluster distinctly from both 6 to 8 and 10 to 12 hour embryos, with far less differentiation among the 2 to 4 hour nuclei than is observed for the older embryos containing differentiated tissues [45].

These findings are not limited to the fly. Studies probing chromatin structure in pluripotent stem cells and mouse blastocysts show that cells in the pluripotent state possess a unique chromatin landscape, marked by a general openness, loosely organized euchromatin, a lack of heterochromatin structure, and reduced repressive histone marks [46,47], a feature we have observed in the early fly embryo [2]. Additionally, it has been shown that this general openness becomes more compact upon differentiation [46,48] and that chromatin accessibility profiles of embryonic stem cells are distinct from precursor and differentiated cell types [49].

The similarity of chromatin profiles in the anterior and posterior halves of the embryo is consistent with our previously described model in which chromatin accessibility at the promoters and enhancers of patterned genes is established in a spatially uniform manner, and not, as is widely believed, by the spatially patterned systems that regulate the activity of these regions.

Work from our lab and others has implicated the ubiquitously active, maternal factor Zelda in this process [2,21–24,50]. The vast majority of enhancers and promoters of genes patterned in the early embryo have binding sites for Zelda and are bound by Zelda prior to mitotic cycle 10 [21]. Our model posits that this early binding of Zelda ensures that these regions remain in an open state as the genome becomes less generally accessible through subsequent mitotic cycles [2,50]. Then, as the patterning systems become active before and during mitotic cycle 14, patterning transcription factors bind to already accessible regions and determine their specific activity.

While Zelda is clearly an important component of this ubiquitous chromatin opening system, there are almost certainly other players, including the ubiquitously expressed trithorax-like/GAGA Factor (or GAF) which plays an important role in establishing accessibility at promoters [22,51–53] and is likely associated with changes in the nucleo-cytoplasmic ratio [50,54].

It is intriguing that, while A-P enhancers were accessible in both embryo halves regardless of where they are active, the magnitude of their accessibility was modestly but significantly skewed in the direction of activity. There are two obvious explanations for this. First, the binding of patterning factors could modify the chromatin effects of Zelda such that enhancers become more open where they are active. Alternatively, the establishment of chromatin accessibility at enhancers showing the most skew could be influenced by a Zelda-independent system with a spatial bias. We believe available data support the latter of these two possibilities.

The strongest accessibility skew is in enhancers with an anterior bias, which as a group are bound highly by Bicoid (BCD), a maternally deposited transcription factor that is expressed in a concentration gradient and specifies head structures [12,55,56], but not bound highly by Zelda or other assayed A-P transcription factors (Fig. 5, Fig. S4). Furthermore, it has recently been shown that there is a class of Bicoid responsive, Zelda independent enhancers whose accessibility is reduced in Bicoid mutant embryos [19], with most of our most strongly skewed enhancers falling into this BCD-responsive, ZLD-independent class.

In conclusion, the data we present here are inconsistent with a model in which chromatin accessibility is a byproduct of activity, and support multiple lines of evidence that the identification of active regions through the establishment of open chromatin in *Drosophila* is largely driven by factors that act in a spatially uniform manner.

## Methods and Materials

### Fly lines

*Drosophila melanogaster* OregonR embryos were collected for 2 hours and aged for 90 minutes on molasses agar plates. Embryos were then dechorionated with 30%-50% bleach solution for three minutes. Embryos were hand staged at 20x magnification at 14°C to be mitotic cycle 14 (NC14) using previously established methods [2].

### Slicing frozen embryos

NC14 embryos were placed in a custom freezing buffer consisting of ATAC-seq lysis buffer [25] without detergent, 5% glycerol, and 1ul of bromoblue dye. Embryos were then taken out of the freezing buffer and placed onto a glass slide which was then put on dry ice for 2-5 minutes. Once embryos were completely frozen, the glass slide was removed and embryos were sliced with a chilled razorblade. Sliced embryo halves were moved to tubes containing ATAC-seq lysis buffer with 0.15mM spermine added to help stabilize chromatin.

### ATAC-seq on frozen embryo halves

Embryo halves were then homogenized using Kimble Kontes Pellet Pestle (cat no. K749521-1590). IGEPal CA-630 was added to a final concentration of 0.1%. After a 10 minute incubation nuclei were spun down and resuspended in water. Twenty halves were added to the transposition reaction containing 25ul of 2x TD buffer (Illumina), and 7.5ul of Tn5 enzyme (Illumina). The reaction was incubated at 37°C for 30 minutes. Transposed DNA was purified using Qiagen Minelute kit. Libraries were then amplified using Phusion (NEB cat no. F531S) and Illumina Nextera index kit (cat no. FC-121-1011). Libraries were then purified with Ampure Beads at a 1.2: 1 beads to sample ratio and sequenced on the Hiseq4000 using 100bp paired end reads. Fragments over 500bp were removed from libraries using a Pippen prep to reduce sequencing bias with the Hiseq4000.

### ATAC-seq data preprocessing and normalization

Fastq files were aligned to *Drosophila* dm3 genome with Bowtie2 using the following parameters -5 5 -3 5 -N 1 -X 2000 --local --very-sensitive-local. Mapping metrics are provided in supplementary table 2. Sam files were then sorted and converted to Bam files using Samtools, only keeping uniquely mapped reads with a MAPQ score of 30 or higher using -q 30. Duplicates were removed with Picard. Bams were then converted to bed files with bedtools and shifted using a custom shell script to reflect a 4bp increase on the plus strand and a 5bp decrease on the minus strand as recommended by [25]. Replicate bed files were merged. Finally shifted bed files were converted into wig files using custom scripts and wig files which were uploaded to the genome browser: https://genome.ucsc.edu/cgi-bin/hgTracks?hgS_doOtherUser=submit&hgS_otherUserName=jhaines&hgS_otherUserSessionName=082917_ATAC%2DseqHalves_paperSession. Merged wig files were normalized to reflect 10 million mapped *Drosophila melanogaster* reads. Anterior, posterior and combined samples were normalized by linear regression to the whole embryo sample not including the y intercept (Fig. S1).

### Peaks

Replicates were merged and peaks were called on the merged bed file using MACS2 with the following parameters: --nomodel --nolambda --keep-dup all --call-summits.

### A-P and D-V patterned enhancer and promoter annotations

A-P and D-V patterned enhancer and gene annotations were compiled from many sources (File S1) [27–39,57]. In order to provide the most accurate promoter annotations possible for our analysis we used RACE, CAGE, and EST data performed in *Drosophila melanogaster* embryos [58] to identify promoters preferentially used by the fly embryo. When there were multiple promoters per gene (as was frequently seen), we chose the promoter that was verified by all three methods, denoted by a “V” in Hoskins et. al. (2011) supplementary file 3. There were three genes that did not have annotated promoters in the Hoskins et al. (2011) dataset that were used in our analysis. Instead these promoter annotations came from the Eukaryotic Promoter Database converted to dm3 annotations [59,60].

In order to further validate our A-P and D-V patterned enhancer and promoter annotations we manually curated *in situ* hybridization images corresponding to 678 genomic regions from multiple sources [28,32,36–38,61–84]. Each region was manually inspected such that only regions with both an *in situ* hybridization image that shows spatially restricted expression as well as moderate accessibility signal (wig signal > 200) were kept for further analysis leaving 253 enhancers and promoters with spatially restricted expression. A report PDF for each region, including in situs, accessibility browser traces, and Z-score and p-value, are found in Supplementary file 2.

### Differential ATAC-seq Analysis

All graphs were made with R scripts (File S3) and Deeptools [85]. Accessibility skew score was calculated by the following equation:

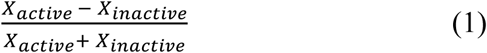

where X_active_ is the wig signal in the half where the region is activating gene expression and X_inactive_ is wig signal in the half where the region is not supposed to activate gene expression. Accessibility skew score measures whether a region is differentially accessible in the expected direction. This score is useful when comparing differential accessibility regardless of which half is favored (for example when comparing accessibility skew at anterior to posteriorly patterned regions).

Positional accessibility skew score provides information about the direction of the skew such that regions that are more accessible in the anterior have a positive positional score while those that are more accessible in the posterior have a negative positional score. Positional accessibility skew score is calculated by the following equation:

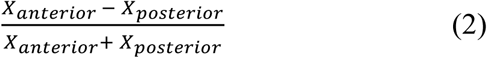

Where X_anterior_ is the wig signal in the anterior sample and X_posterior_ is the wig signal in the posterior sample. Significance for each region was determined by computationally matching each region to a random region that has the same total normalized wig score (Fig. S5). Accessibility skew score was calculated for each random region (termed RandSkewScore). These scores were distributed normally and allowed for determining a Z-score for each region of interest (Z_ROI_) by the following equation:

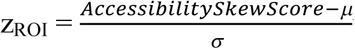where μ is the mean of the random region distribution and μ is the standard deviation of the random region distribution. 2 tailed P-values were then calculated from the Z score.

### ChIP-seq data analysis

Wig files from previously published ChIP-seq data were obtained for Kruppel, Hunchback, Giant, Knirps, Caudal, Bicoid [40], and Zelda data from stage 3,4, and 5 embryos [21]. Wig files were normalized by the mean signal for each sample, assuming that the mean signal over the entire genome is similar to that of background. This normalization essentially transforms the data into deviations from the mean such that signal from different experiments can be compared to each other. Wig signal around regions of interest was determined and graphed in R (File S3). For the heatmaps, normalized wig signal was summed over 3kb windows around regions of interest for each factor before being scaled such that the region with the highest value is equal to 1 and the lowest to 0 for each factor.

Data is currently being submitted to NCBI GEO but can also be accessed via http://eisenlab.org/aphalves/.

## Acknowledgments

This research was supported by the MBE’s Investigator Award from the Howard Hughes Medical Institute and JEH’s Graduate Research Fellowship from the National Science Foundation and NHGRI “T32” training grant in Genomics and Computational Biology. The authors would like to thank members of MBE’s research group for comments on an earlier draft of this manuscript. Special thanks to Mawupedzro K Tekpa for help creating the *in situ* hybridization database and Xiao-Yong Li, Michael Stadler, Steven Kuntz, and Ashley Albright, and for help optimizing ATAC-seq on frozen embryo slices, data analysis, and figure design.

---

**Figure S1.**
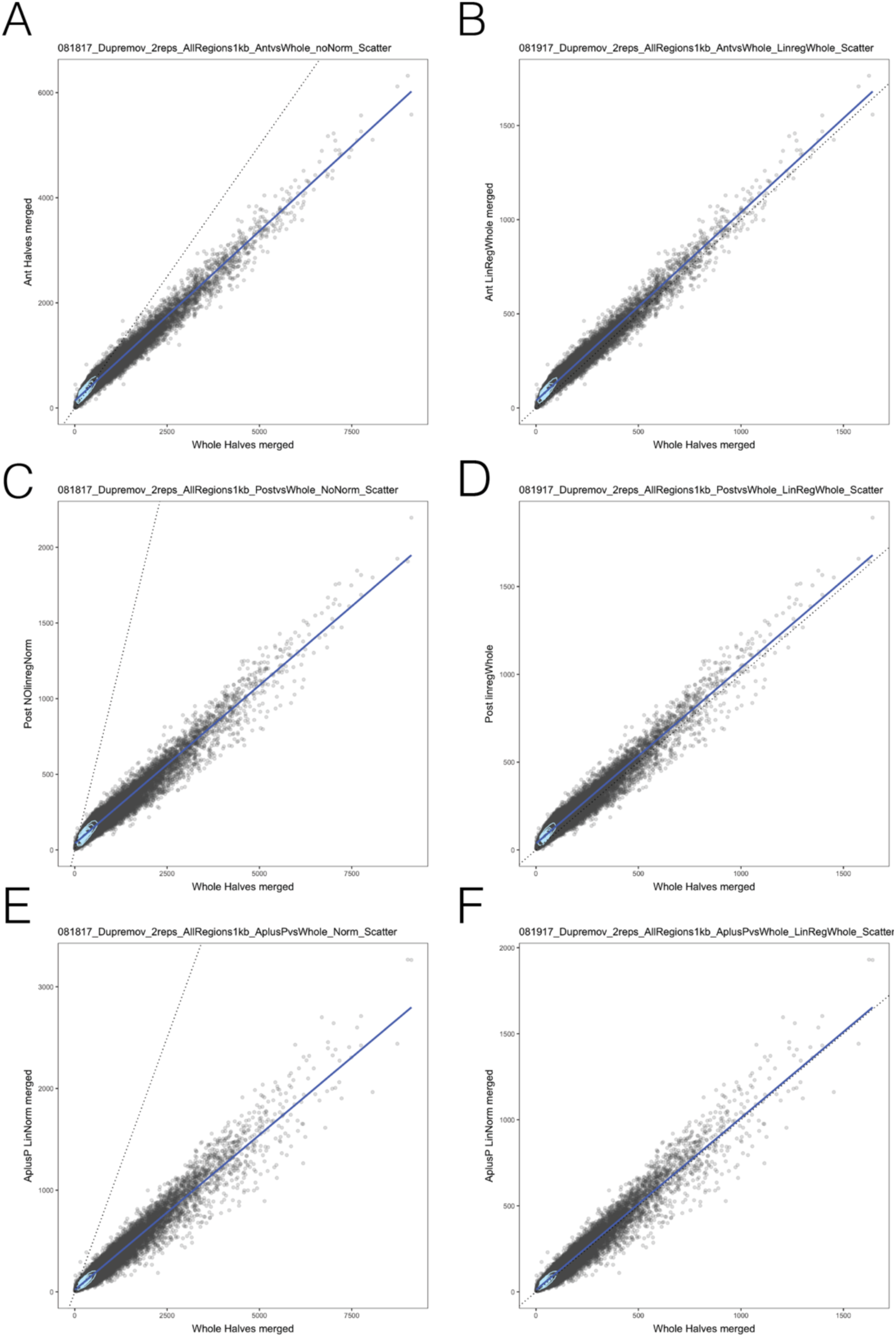
Linear regression normalization. Scatterplots showing 1kb genomic bins (gray) with 1D density plot (light blue) indicating areas of increased point density. X=Y line indicated by dashed line. Linear regression line is indicated by a solid dark blue line. Anterior, posterior, and combined halves were normalized to whole samples. (A,C,E) Scatterplots show data before normalization (B,D,F) Scatterplots show data after normalization.

**Figure S2.**
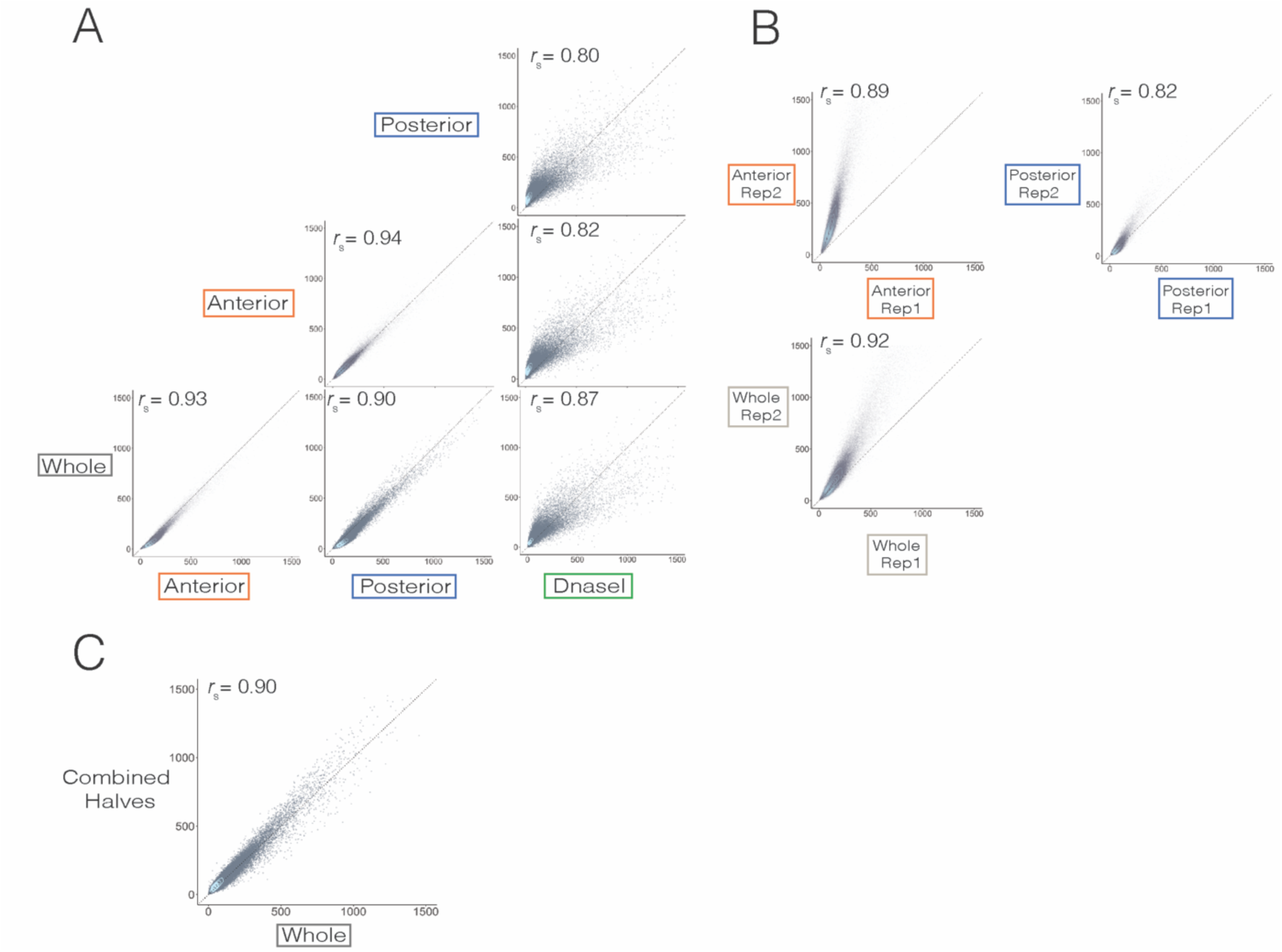
ATAC-seq on embryo halves correlates highly with DnaseI and whole embryos. (A) Scatterplots showing merged normalized wig signal values from ATAC-seq experiments performed on anterior halves, posterior halves, and whole embryos compared with DNaseI data from stage 5 *Drosophila melanogaster* embryos (Thomas et al. 2011) binned into 1kb regions. Spearman correlation coefficient is shown above each plot. The line X=Y is shown as a dotted line. (B) Scatterplots showing non-normalized wig signal values for anterior halves, posterior halves, and whole replicates. (C) Normalized wig signal from ATAC-seq data for the combined anterior and posterior halves sample compared to the merged whole embryo sample.

**Figure S3.**
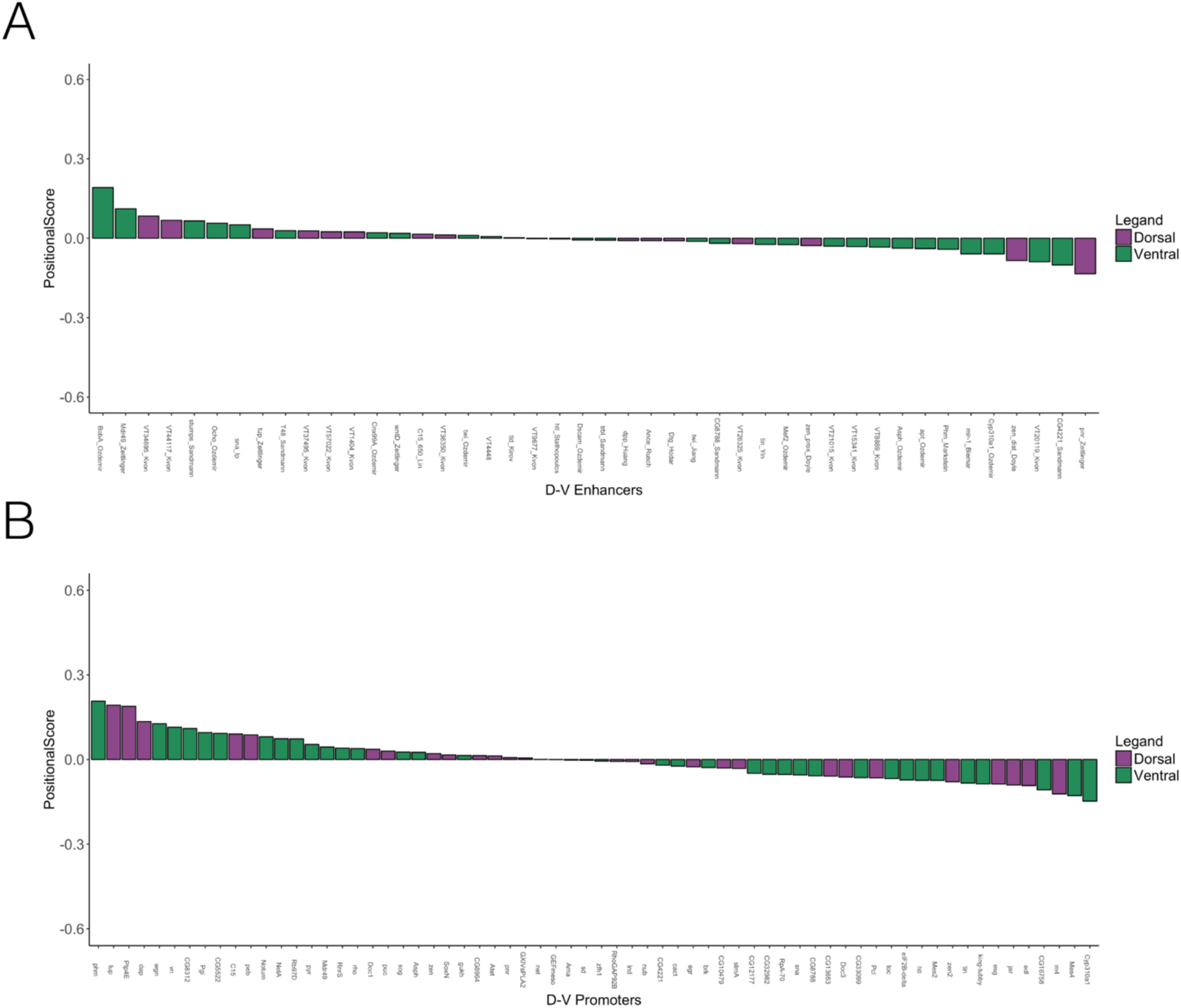
D-V patterned enhancers and promoters are similarly accessible in both halves. Bargraphs showing positional score calculated for dorsal (purple) and ventral (green) patterned enhancers and promoters. The enhancer or promoter names are below the graph. (A) D-V enhancers (B) D-V promoters

**Figure S4.**
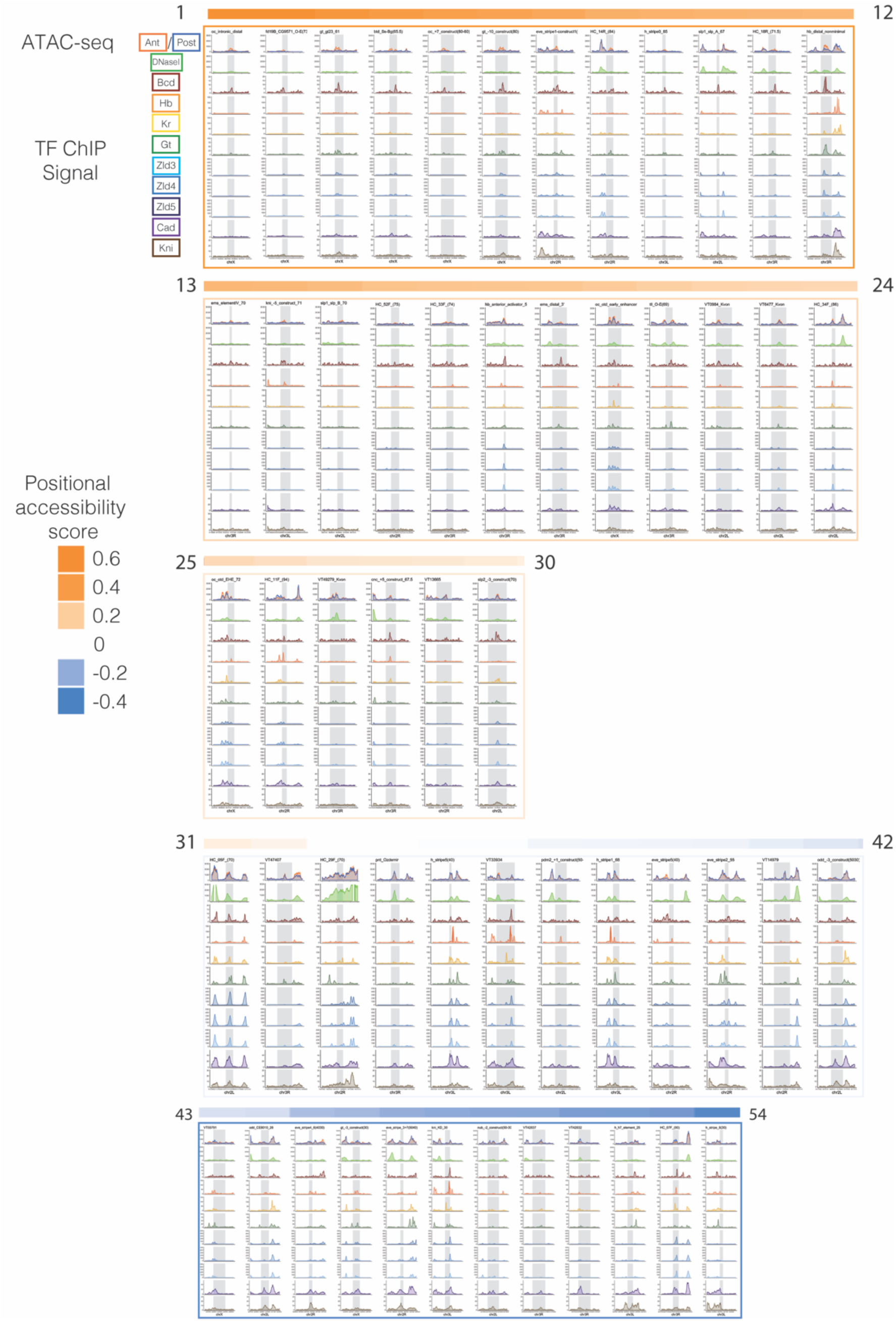
A-P patterned transcription factor binding at similarly and differentially accessible A-P patterned enhancers. A-P patterned enhancers from Fig3 are ordered from by positional accessibility skew score (positional score is indicated by the colored bar above each panel – orange indicates more differentially accessible in the anterior, blue indicates more differentially accessible in the posterior, and white is similarly accessible in both halves. Each panel consists of normalized wig signal in a 3kb window around each enhancer (the actual enhancer region is denoted by a gray rectangle). The first panel shows normalized, merged, ATAC-seq signal in the anterior (orange) and posterior (blue) halves. The second panel shows the DNAseI signal from stage 5 embryos (Thomas et al. 2011) in green. The third through 11^th^ panels are normalized wig signal from ChIP-seq experiments of the following proteins: Bicoid (red), Hunchback (orange), Kruppel (yellow), Giant (green), Zelda from stage 3,4,and 5 embryos (light, medium, and dark blue), Caudal (purple), and Knirps (brown). The name of the enhancer is above each panel.

**Figure S5.**
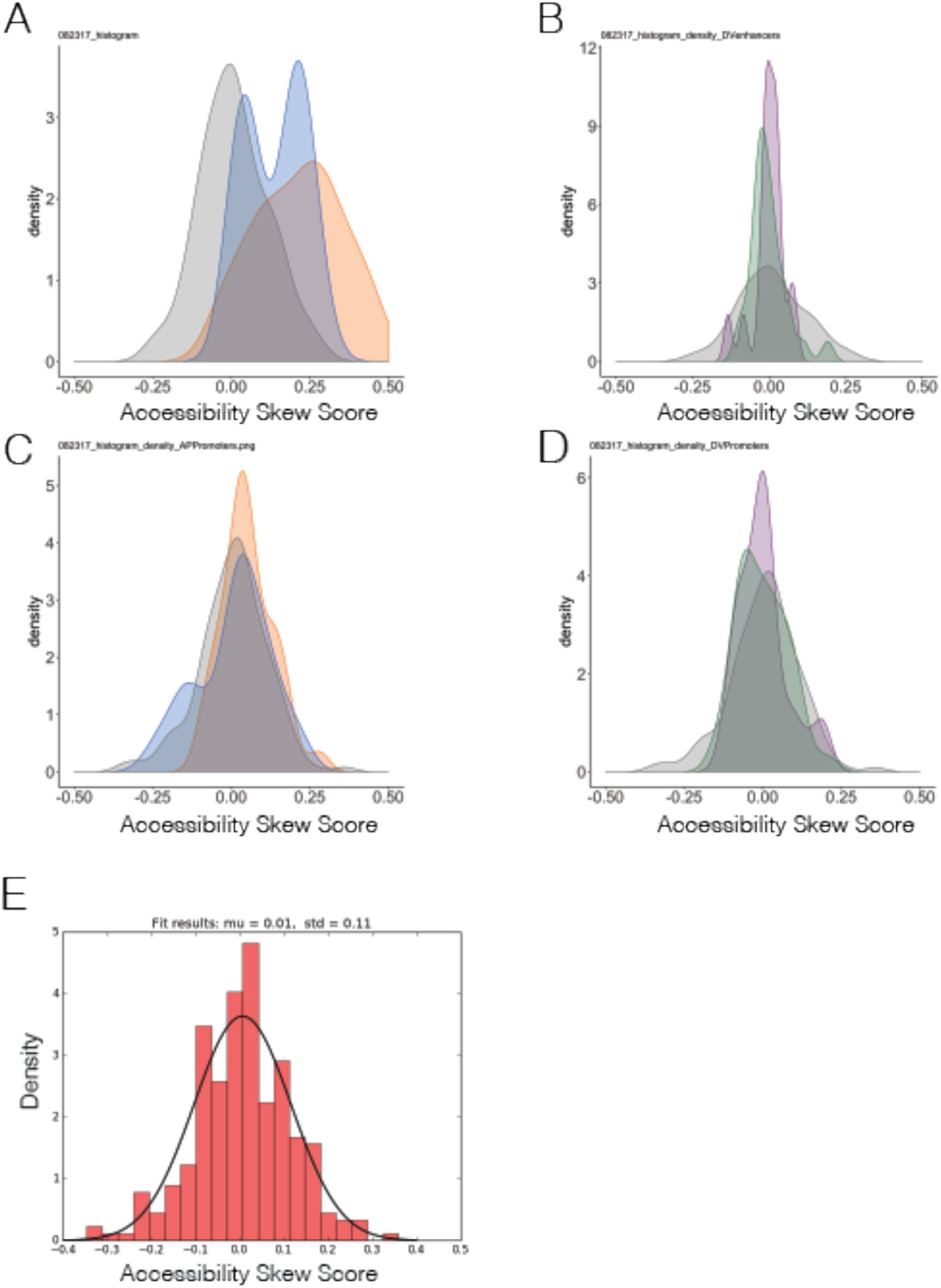
Accessibility skew scores of enhancers and promoters compared to random regions. Histograms showing distribution of accessibility skew scores of random regions compared to (A) A-P patterned enhancers (B) D-V patterned enhancers (C) A-P patterned promoters (D) D-V patterned promoters. Anterior is orange, Posterior is blue, Dorsal is purple, Ventral is green. (E) Histogram showing the distribution of random regions with the fitted normal curve in black. Mu and Std from the normal curve is shown above the graph.

**Table S1.** ANOVA Comparisons between A-P and D-V enhancers and promoters.

| Anova Comparisons Between A-P and D-V enhancers and promoters |  |  |  |  |
| --- | --- | --- | --- | --- |
| Enhancers | diff | lwr | upr | p adj |
| Posterior-Anterior | -0.077057528 | -0.16142451 | 0.007309454 | 0.091512659 |
| Dorsal-Anterior | -0.214728907 | -0.300734236 | -0.128723578 | 7.81E-10 |
| Ventral-Anterior | -0.216575473 | -0.290980158 | -0.142170789 | 1.03E-12 |
| Random-Anterior | -0.205945352 | -0.262255675 | -0.149635029 | 1.10E-13 |
| Dorsal-Posterior | -0.137671379 | -0.236512224 | -0.038830535 | 0.001569157 |
| Ventral-Posterior | -0.139517945 | -0.228448553 | -0.050587338 | 0.000237485 |
| Random-Posterior | -0.128887824 | -0.203342161 | -0.054433486 | 3.54E-05 |
| Ventral-Dorsal | -0.001846566 | -0.092332929 | 0.088639797 | 0.999997724 |
| Random-Dorsal | 0.008783556 | -0.067522258 | 0.08508937 | 0.997800125 |
| Random-Ventral | 0.010630121 | -0.052312056 | 0.073572299 | 0.990345927 |
| Promoters | diff | lwr | upr | p adj |
| Posterior-Anterior | -0.043074207 | -0.107594479 | 0.021446065 | 0.356680419 |
| Dorsal-Anterior | -0.050179092 | -0.12052014 | 0.020161956 | 0.289167405 |
| Ventral-Anterior | -0.0531619 | -0.11808839 | 0.01176459 | 0.165304979 |
| Random-Anterior | -0.046359571 | -0.098341591 | 0.005622449 | 0.105683775 |
| Dorsal-Posterior | -0.007104885 | -0.07625218 | 0.06204241 | 0.998608073 |
| Ventral-Posterior | -0.010087693 | -0.073718931 | 0.053543546 | 0.992507483 |
| Random-Posterior | -0.003285364 | -0.053640262 | 0.047069534 | 0.999768593 |
| Ventral-Dorsal | -0.002982808 | -0.072509292 | 0.066543676 | 0.999956364 |
| Random-Dorsal | 0.003819521 | -0.053805242 | 0.061444283 | 0.999753624 |
| Random-Ventral | 0.006802329 | -0.044072019 | 0.057676677 | 0.996110657 |

**Table S2.** ATAC-seq halves mapping metrics.

| Sample | All reads | % Overall Alignment Rate (including multiple) | Final Read number (filtered for quality score + dup removed) |
| --- | --- | --- | --- |
| ME_JH_112315_1105-ATACSlice2-01-20a | 36984564 | 92.69 | 14622650 |
| ME_JH_112315_1105-ATACSlice2-02-20p | 19383698 | 96.75 | 8850316 |
| ME_JH_112315_1105-ATACSlice2-5-1A | 38641395 | 91.54 | 9996838 |
| ME_JH_112315_1105-ATACSlice2-6-1p | 38796161 | 72.82 | 2821012 |
| ME_JH_112315_1105-ATACSlice2-7-10whole | 47244276 | 97.68 | 23148054 |
| ME_JH_112315_1109-ATACSlice03-01-20Ant | 50087614 | 95 | 40364768 |
| ME_JH_112315_1109-ATACSlice03-02-20Post | 34457240 | 38.15 | 8753933 |
| ME_JH_112315_1109-ATACSlice03-03-1plus2 | 42215974 | 54.63 | 16630998 |
| ME_JH_112315_1109-ATACSlice03-04-10whole | 43706348 | 96.69 | 32344883 |

